# Cyclophilin A Facilitates HIV-1 DNA Integration

**DOI:** 10.1101/2024.06.15.599180

**Authors:** Adrian Padron, Richa Dwivedi, Rajasree Chakraborty, Prem Prakash, Kyusik Kim, Jiong Shi, Jinwoo Ahn, Jui Pandhare, Jeremy Luban, Christopher Aiken, Muthukumar Balasubramaniam, Chandravanu Dash

**Affiliations:** Center for AIDS Health Disparities Research; Department of Microbiology, Immunology, and Physiology; School of Graduate Studies; Department of Biochemistry, Cancer Biology, Pharmacology, and Neuroscience Meharry Medical College, Nashville, TN; Program in Molecular Medicine University of Massachusetts Medical School, Worcester, MA; Department of Pathology, Microbiology, and Immunology Vanderbilt University Medical Center, Nashville, TN; Department of Structural Biology University of Pittsburgh, Pittsburgh, PA

**Keywords:** Cyclophilin A (CypA), Human immunodeficiency virus (HIV), capsid, reverse transcription nuclear entry, integration

## Abstract

Cyclophilin A (CypA) promotes HIV-1 infection by facilitating reverse transcription, nuclear entry and by countering the antiviral activity of TRIM5α. These multifunctional roles of CypA are driven by its binding to the viral capsid. Interestingly, recent studies suggest that the HIV-1 capsid lattice enters the nucleus of an infected cell and uncoats just before integration. Therefore, we tested whether CypA-capsid interaction regulates post-nuclear entry steps of infection, particularly integration. First, we challenged CypA-expressing (CypA^+/+^) and CypA-depleted (CypA^-/-^) cells with HIV-1 particles and quantified the resulting levels of provirus. Surprisingly, CypA-depletion significantly reduced integration, an effect that was independent of CypA’s effect on reverse transcription, nuclear entry, and the presence or absence of TRIM5α. Additionally, cyclosporin A, an inhibitor that disrupts CypA-capsid binding, inhibited HIV-1 integration in CypA^+/+^ cells but not in CypA^-/-^ cells. Accordingly, HIV-1 capsid mutants (G89V and P90A) deficient in CypA binding were also blocked at integration in CypA^+/+^ cells but not in CypA^-/-^ cells. Then, to understand the mechanism, we assessed the integration activity of HIV-1 preintegration complexes (PICs) extracted from infected cells. The PICs from CypA^-/-^ cells had lower activity *in vitro* compared to those from CypA^+/+^ cells. PICs from cells depleted for CypA and TRIM5α also had lower activity, suggesting that CypA’s effect on PIC activity is independent of TRIM5α. Finally, addition of CypA protein significantly stimulated the integration activity of PICs extracted from both CypA^+/+^ and CypA^-/-^cells. Collectively, these results suggest that CypA promotes HIV-1 integration, a previously unknown role of this host factor.

**Importance:** HIV-1 capsid interaction with host cellular factors is essential for establishing a productive infection. However, the molecular details of such virus-host interactions are not fully understood. Cyclophilin A (CypA) is the first host protein identified to specifically bind to the HIV-1 capsid. Now it is established that CypA promotes reverse transcription and nuclear entry steps of HIV-1 infection. In this report, we show that CypA promotes HIV-1 integration by binding to the viral capsid. Specifically, our results demonstrate that CypA promotes HIV-1 integration by stimulating the activity of the viral preintegration complex and identifies a novel role of CypA during HIV-1 infection. This new knowledge is important because recent reports suggest that an operationally intact HIV-1 capsid enters the nucleus of an infected cell.

## Introduction

Approximately seventy-nine million people have been infected with the human immunodeficiency virus type 1 (HIV-1) worldwide [1]. Unfortunately, an estimated forty million of these individuals have died and currently over thirty-nine million people are carrying the virus [1]. HIV-1 primarily infects immune cells such as T-lymphocytes, monocytes, and macrophages that that express CD4 receptor and CCR5 or CXCR4 co-receptors [2–8]. The major consequence of HIV-1 infection is the steady decline of CD^4+^ T cells [9–12], resulting in the loss of immune control and the development of immunodeficiency [10, 13]. Particularly, immunodeficiency is caused by T-cell dysfunction, defects in both number and function of antigen-presenting cells, thymic dysfunction, lymphoid destruction, and stem cell dysfunction [10–12]. Without antiretroviral therapy, these HIV-1 infection-associated immunodeficiencies lead to the deadly form of the disease called acquired immunodeficiency syndrome (AIDS) [11, 14].

Structurally, HIV-1 consists of an outer lipid membrane that surrounds a conical capsid [15]. The capsid shell consists off ∼200-250 hexamers and 12 pentamers of the viral capsid (CA/p24) protein [16]. The core of the capsid contains a number of viral and host factors as well as two copies of the viral single stranded RNA genome [17–19]. After fusion of viral membrane with the target cell membrane, the viral capsid is released into the cytoplasm of the target cell and is trafficked towards the nuclear pore complex (NPC) [20–22]. Concomitantly, the core-associated reverse transcription complex (RTC) containing the viral reverse transcriptase (RT) begins to synthesize a double stranded DNA from the viral RNA genome [21, 23]. The newly synthesized viral DNA is transported into the host cell nucleus through the NPC. Then, the preintegration complex (PIC) containing the viral integrase (IN), and other factors, inserts the viral DNA into actively transcribing genes of the host chromosomes [24]. HIV-1 DNA integration step establishes a provirus in the host genome that serves as a permanent genetic element for the production of progeny virions [24]. However, these early steps of HIV-1 infection are critically dependent on binding of host factors/proteins to the viral capsid [25, 26]. For instance, trafficking of the viral capsid to the NPC is facilitated by the kinesin adapter protein FEZ1 [27, 28]. Additionally, host proteins such as Sec24C, Nucleoporins (Nups: e.g., Nup358 and Nup153), Transportins (e.g., TNPO1 and TNPO3), Bicaudal D2 (BICD2) and cleavage and polyadenylation specificity factor 6 (CPSF6) coordinate viral nuclear entry [29–37]. CPSF6 also supports integration targeting into transcriptionally active regions of the human chromosomes [38–42]. Cyclophilin A (CypA) is another important CA-binding host factor that plays an expansive role during the early steps of HIV-1 infection.

Human CypA is an abundantly expressed cytosolic protein and is encoded by the *PPIA* gene located in chromosome 7 [43, 44]. CypA belongs to the cyclophilin family of proteins that are structurally and functionally conserved in prokaryotes and eukaryotes [45, 46]. Structurally, CypA is an 18 kDa protein containing a cyclophilin-like domain (CLD) [47] with two alpha-helices and a beta sheet [48, 49]. Functionally, CypA possesses the peptidyl/prolyl *cis-trans* isomerase (PPIase) activity [50] that isomerizes the peptide bond preceding proline residues [51–53]. The PPIase activity of CypA is required for protein folding, protein trafficking/molecular chaperoning, cell signaling, and T cell activation [46, 51, 54–56]. Therefore, CypA has been implicated in a number of physiological and disease conditions including infections of viruses such as influenza, hepatitis C virus, coronavirus, and HIV [57–59].

CypA’s role in HIV-1 infection was first demonstrated in a study where CypA was shown to bind to the HIV-1 Gag polyprotein [60]. Specifically, CypA was found to bind to the proline-rich loop (aka CypA-binding loop) located in the N-terminal domain (NTD) of the CA region of the Gag [61–63]. It is now established that CypA-CA binding promotes multiple steps of HIV-1 infection starting from reverse transcription to integration targeting. For instance, HIV-1 particles assembled in the presence of a CypA-inhibitor are less infectious [64] and the lack of virion-associated CypA impairs viral DNA synthesis [65, 66]. Similarly, point mutations P90A or G89V/A in the CypA binding loop of CA reduce viral DNA synthesis [65–70]. Although the precise mechanism is not fully understood, the positive effects of CypA on HIV-1 reverse transcription have been linked to its ability to stabilize the viral capsid, potentially by blocking the binding and restriction activity of the host restriction factor TRIM5α [70–76]. Similarly, CypA coordinates binding of Nups to the viral capsid to regulate HIV-1 nuclear entry [32, 77–79]. Accordingly, when CypA-CA interaction is disrupted, utilization of these Nups is altered for the nuclear entry of the virus [32, 77, 80–82]. Interestingly, CypA-CA interaction also regulates the rate at which HIV-1 replication complex docks at the NPC [39, 83]. However, CypA’s effect on HIV-1 nuclear entry is cell type dependent [29] and CypA could inhibit nuclear entry of HIV-1 in old world monkey cells [84]. Furthermore, CA mutant (G89V, P90A) viruses defective for CypA binding are also resistant to Myxovirus resistance B (MxB)-mediated inhibition of HIV-1 nuclear entry [85–87]. Collectively, the current model suggests that CypA-CA interaction is important for the import, docking and entry of HIV-1 replication complex through the NPC into the nucleus of an infected cell. Finally, there is indirect evidence that CypA-CA interaction could influence post-nuclear entry steps such as HIV-1 integration targeting [32]. For example, disruption of CypA-CA binding has been reported to increase targeting of HIV-1 integration into gene-dense regions [32]. This contrasts with depletion of CPSF6, where integration targeting is directed away from gene-dense regions [32, 38, 42]. However, the mechanism underlying the surprising effects of CypA on integration targeting remains largely unknown, most likely because a direct role of CypA in post-nuclear entry steps of HIV-1 infection has not been studied.

Human CypA is predominantly cytosolic in its cellular distribution, thus a nuclear role of this host factor in HIV-1 permissive cells either in normal cellular function or diseased conditions has remained elusive. Nevertheless, a recent study detected CypA in the nucleus of human monocyte-derived macrophages (MDMs) [83]. CypA has also been reported to localize to the nucleus of Jurkat T-cells during the completion of cytokinesis [88] and upon stimulation of cells with stressors to play an anti-apoptotic role [89]. Coincidently, recent reports also demonstrate that intact or near-intact HIV-1 capsid can be observed inside the nuclear pores and inside the nucleus of infected cells [90–92]. Most importantly, we have reported that CypA regulates the integration step of HIV-1 CA mutant (R264K) that evades the antiviral effects of the cytotoxic T lymphocytes (CTLs) [93]. Accordingly, the compensatory CA mutation S173A restored the integration and infectivity defect of the R264K mutant. These results suggest either a direct or a functional role of CypA in post-nuclear entry steps of HIV-1 infection.

The premise for a nuclear role of CypA during HIV-1 infection is also derived by our study demonstrating that HIV-1 CA controls viral DNA integration [94]. We have reported that the anti-HIV-1 activity of the CA-inhibitor PF74 (PF-3450074) maps to the block at the integration step [94]. PF74 binds to a pocket at the NTD–CTD inter-subunit interface of CA and exhibits a concentration-dependent anti-viral activity [33, 95–98]. At higher concentrations (>2 µM), PF74 inhibits HIV-1 reverse transcription and nuclear entry [33, 96, 99], presumably by triggering premature capsid uncoating [98–100]. Interestingly, we found that at 2 µM, PF74 blocked proviral integration and this reduction correlated quantitatively with antiviral activity. Furthermore, our biochemical studies revealed that PF74 inhibits HIV-1 PIC-associated viral DNA integration *in vitro*. This inhibitory effect is CA-specific since PF74 did not reduce the integration activity of PICs of the PF74-resistant (5Mut) HIV-1. Additionally, partial purification of PICs and immunoprecipitation assay, demonstrated that CA is associated with the functional PICs. Since PF74’s anti-viral effect is dependent on its binding to CA, these results supported a direct role of CA in HIV-1 integration. Therefore, in this study we predicted that CypA, as a CA-binding host factor, could play a direct role in HIV-1 integration. To test this prediction, we combined molecular virology, genetic, pharmacologic and biochemical approaches and describe that CypA promotes HIV-1 integration. Notably, we found that CypA’s effect on HIV-1 integration was independent of its effect on reverse transcription and nuclear entry. Our results also suggest that the effects of CypA on HIV-1 integration is dependent on its ability to bind to the viral capsid. Additionally, our study provide evidence that CypA’s effect on HIV-1 integration is not dependent on TRIM5α. Finally, our biochemical studies of HIV-1 PICs indicated that CypA promotes HIV-1 integration by stimulating PIC-mediated viral DNA integration activity. Collectively, these observations suggest a direct and functional role of CypA in HIV-1 DNA integration, a previously unknown role of this host factor in post-nuclear entry steps of infection.

## Materials and Methods

### Chemicals

The antiretroviral compounds Efavirenz and Raltegravir were obtained through BEI Resources (NIH HIV Reagents Program; Manassas, VA) and prepared as previously described [94]. Cyclosporine A was purchased from MedChemExpress (Monmouth Junction, NJ) and reconstituted in DMSO as per manufacturer’s instructions. The recombinant cyclophilin

### Cell culture

TZM-bl and HEK293T cell lines were obtained from the American Type Cell Culture Collection (Manassas, VA). The CD4+ T lymphocyte Jurkat (parental CypA^+/+^) cell line and its cyclophilin A knock-out derivative (CypA^-/-^) cell line were obtained from BEI Resources/NIH HIV Reagents Program (Germantown, MD). The CypA^-/-^/TRIM5α^-/-^ double knockoutJurkat cell line was a kind gift from Jeremy Luban (University of Massachusetts, Worcester MA). The TZM-bl and HEK293T cells were cultured in Dulbecco’s Modified Eagle’s medium (ThermoFisher Scientific) containing 10% heat-inactivated fetal bovine serum (FBS), penicillin (50 IU/mL), streptomycin (50 µg/mL), and L-glutamine (2 mM). All the Jurkat cell lines were cultured in RPMI 1640 media containing 10% heat-inactivated FBS, penicillin (50 IU/mL), streptomycin (50 µg/mL), and L-glutamine (2 mM). All cell lines were cultured at 37°C with 5% CO_2_.

### Viral stock preparation

The viruses utilized in this study were generated from the infections molecular clone pNL4-3 [101]; and its CA mutant derivatives (G89V and P90A). The plasmid was purchased from BEI Resources. The proviral molecular plasmids, pNL4-3, pNL4.3 CA mutants G89A and P90A were used to generate HIV-1 viral particles. Infectivity and viral DNA quantification experiments were performed with VSV-G pseudotyped NL4-3 virions which were generated from the pNL4-3 plasmid that contains a frameshift in the *env* gene and the GFP-coding sequence inserted in place of the *nef* gene [102]. All virus stocks were prepared as previously described [94]. Briefly, 5 x 10^6^ 293T cells were seeded in 10 cm cell culture plates and cultured overnight. Next day, cells in each culture plate were transfected with 10µg of pNL4-3 plasmid DNA complexed with 50μL PEI reagent (Sigma-Aldrich catalog # 764604), and the culture media was changed 16 hours post-transfection. Forty-eight hours later, the virus-containing culture media was collected, subjected to low-speed centrifugation, and the supernatant was passed through 0.45µm filter. The filtrate was treated with DNase I (ThermoFisher Scientific catalog #AM2238) at a final concentration of 20μg/mL at 37°C. The viral stocks were stored in −80°C freezer until assayed for infectivity. Subsequently, ELISA was used to assay the virus preparations for p24 amounts using anti-p24 antibody as previously described [94].

### Infectivity assays

The infectivity of all viruses was determined using the TZM-bl cell line-based luciferase reporter assay (Reference). Briefly, 5 x 10^4^ TZM-bl cells were seeded in each well of 24-well cell culture plates and cultured for 24 hours. The cells were inoculated with appropriate virus inoculum in the presence of polybrene (Sigma-Aldrich catalog # TR-1003) and cultured for 2 hours in the 37°C/5% CO_2_ incubator. Then, the inoculum medium was removed and replaced with new complete media and the cells were cultured for 48 hours in 37°C/5% CO_2_ incubator. Cells were lysed Passive Lysis Buffer (Promega, catalog # E1941) and the luciferase activity was determined using BrightGlo^TM^ Luciferase Assay System (Promega, catalog # 2610) in a microplate reader (BioTek Synergy).

### Western blot

Cells were lysed in radioimmunoprecipitation buffer (Sigma-Aldrich, catalog #R0278) and 20μg of each cell lysate was resolved by 4-12% SDS-PAGE, transferred onto nitrocellulose membrane, and probed with anti-cyclophilin A antibody (Santa Cruz Biotechnology sc-134310), anti-TRIM5α antibody (Cell Signaling Technology #14326), and anti-beta actin (sc-517582) antibody.

### Single-cycle infection assay

1×10^5^ Jurkat cells in each well of a 96-well flat-bottom cell culture plate were spinoculated (480 x g) with 250ng of pseudotyped HIV-1.GFP (Reference) in the absence or presence of cyclosporine A (10μM) for 2 hours at 25°C and were then cultured for 24 hours in 37°C/5% CO_2_ incubator. The cells were then fixed using BD Cytofix/Cytoperm (BD Biosciences) and were assayed for GFP expression by FACS (Cytek Biosciences Guava EasyCyte 5HT cytometer).

### Total DNA isolation for qPCR assays

To isolate total DNA from infected cells, 1×10^6^ Jurkat (parental CypA^+/+^, CypA^-/-^, and CypA^-/-^/TRIM5α^-/-^) cells in each well of a 48-well cell culture plate were inoculated with 2.5μg of pseudotyped HIV-1.GFP and cultured in 37°C/5% CO_2_ incubator. Twenty-four hours post-inoculation, 1×10^4^ cells from each of the inoculated cell samples were fixed and assayed for GFP expression by FACS as a readout of viral infectivity. Then, the remaining inoculated cells were harvested for DNA extraction. Total DNA was extracted from the remaining inoculated cells Quick-DNA miniprep kit (Zymo Research).

### qPCR for measuring the reverse transcription products and 2-LTR circles

qPCR was used to measure the reverse transcription products and 2-LTR circles. The late reverse transcription products were quantified by using SYBR green-based qPCR, and the 2-LTR circles were quantified by using TaqMan probe-based qPCR. The SYBR green-based qPCR mix contained 1X iTaq Universal SYBR Green Supermix (Bio-Rad), 300nM concentrations (each) of forward primer (5’-TGTGTGCCCGTCTGTTGTGT-3’) and reverse primer (5′-GAGTCCTGCGTCGAGAGAGC-3’), and 100 ng of the total DNA from infected cells. The TaqMan probe-based qPCR mix contained 1X iTaq Universal Probe Supermix (Bio-Rad), 300nM (each) of forward primer (5’-AACTAGGGAACCCACTGCTTAAG-3’) and reverse primer (5’-TCCACAGATCAAGGATATCTTGTC-3’), 100nM of the TaqMan probe (5’-[FAM] ACACTACTTGAAGCACTCAAGGCAAGCTTT-[TAMRA]-3’), and 100ng of total DNA from infected cells. The cycling conditions for SYBR green-based qPCR included an initial incubation at 95°C for 3 minutes, followed by 39 cycles of amplification and acquisition at 94°C for 15s, 58°C for 30s, and 72°C for 30s. For the SYBR green-based qPCR, the thermal profile for melting-curve analysis was obtained by holding the sample at 65°C for 31 s, followed by a linear ramp in temperature from 65 to 95°C with a ramp rate of 0.5°C/s and acquisition at 0.5°C intervals. The TaqMan probe-based qPCR included an initial incubation at 95°C for 3 minutes, followed by 39 cycles of amplification and acquisition at 94°C for 15s, 58°C for 30s, and 72°C for 30s. During qPCR of the samples, a standard curve was generated in parallel and under same conditions using 10-fold serial dilutions of known copy numbers (10^0^ to 10^8^) of the HIV-1.GFP plasmid (for late RT products) or the p2LTR plasmid containing the 2-LTR junction sequence (for 2-LTR circles). CFX Manager software (Bio-Rad) was used to analyze the data and determine the copy numbers of the late RT products and the 2-LTR circles by plotting the data against the respective standard curves.

### Alu-gag nested qPCR for measuring HIV-1 proviral integration

To measure integrated HIV-1 proviral DNA in infected T cells, a nested PCR method involving a first round endpoint PCR with primers designed to amplify only the junctions of the chromosomally integrated viral DNA but not any unintegrated viral DNA, followed by a second round of qPCR with primers that amplify viral LTR-specific sequences present in the first round PCR amplicons, was used with some modifications. Briefly, the first-round PCR was performed in a final volume of 50μL containing 100ng of total DNA from infected parental and knockout Jurkat cells, 1X GoTaq reaction buffer (Promega), deoxy-nucleoside triphosphate (dNTP) nucleotide mix containing 200µM concentrations of each nucleotide (Promega), 500nM (each) of primers that target the host chromosomal Alu repeat sequence (5’-GCCTCCCAAAGTGCTGGGATTACAG-3’) and HIV-1 Gag sequence (5’-GTTCCTGCTATGTCACTTCC-3’), and 1.25U of GoTaq DNA polymerase (Promega). The thermocycling conditions included an initial incubation at 95°C for 5 minutes; followed by 23 cycles of amplification at 94°C for 30s, 50°C for 30s, 72°C for 4 minutes and a final incubation at 72°C for 10 min. The second-round qPCR mix contained a 1/10 volume of the first-round PCR products as the template DNA, 1X iTaq Universal Probe Supermix (Bio-Rad), 300nM (each) of the viral LTR-specific primers that target the R region (5’-TCTGGCTAACTAGGGAACCCA-3’) and the U5 region (5’-CTGACTAAAAGGGTCTGAGG-3’), and a 100nM concentration of TaqMan probe (5’-[6-FAM]-TTAAGCCTCAATAAAGCTTGCCTTGAGTGC-[TAMRA]-3’). The qPCR cycling conditions included an initial incubation at 95°C for 3 minutes, followed by 39 cycles of amplification and acquisition at 94°C for 15s and 58°C for 30s, and a final incubation at 72°C for 30s. During the second round qPCR of the samples, a standard curve was generated in parallel and under same conditions using 10-fold serial dilutions of known copy numbers (10^0^ to 10^8^) of the of the HIV-1.GFP plasmid. Data were analyzed using CFX Manager software (Bio-Rad), and integrated viral DNA copy numbers were determined by plotting the qPCR data against the standard curve.

### Extraction of PICs

The HIV-1 PICs were extracted using a published protocol (Reference). Briefly, parental CypA^+/+^, CypA^-/-^, and CypA^-/-^/TRIM5α^-/-^ Jurkat cells (80×106) were acutely spinocualted (480 x g) with NL4-3 HIV-1 (30mL high titer virus) for 2 hours at at 25°C followed by culturing in 37°C/5% CO_2_ incubator for 5 hours. The cells were washed twice with Buffer K^-/-^ (150mM KCl, 20mM HEPES, 5mM MgCl_2_, pH 7.6) and lysed in Buffer K^+/+^ [Buffer K^-/-^ +digitonin (0.025%), 1mM DTT and 20μg/mL aprotinin]. The supernatant containing the cytoplasmic PICs was treated with RNase A and supplemented with sucrose. The PIC preparations were aliquoted, flash frozen, and stored at −80°C.

### *In vitro* integration assay

The *in vitro* integration assays using cytoplasmic PICs were performed as described previously (Reference). Briefly, the *in vitro* integration reactions were assembled by adding 600ng of φX174 2.0kb linear target DNA to PIC (200μL) aliquots and the reactions were incubated at 37°C for 45min. The integration reaction was stopped and deproteinized by adding SDS, EDTA, and proteinase K to final concentrations of 0.5%, 8 mM, and 0.5 mg/mL, respectively, followed by overnight incubation at 56°C (∼16 hours). Total DNA was isolated and purified by phenol/chloroform extraction followed by ethanol precipitation.

### Measurement of PIC-associated viral DNA integration

The integration events from the *in vitro* integration reactions were quantified as copy numbers by using a previously reported nested qPCR-based approach (Reference). Briefly, the DNA isolated from the IVI reactions were used as templates in the first-round end-point PCR that included one-tenth volume of the purified DNA product from the integration reaction, 500nM concentrations (each) of primers targeting the substrate DNA (5’-CACTGACCCTCAGCAATCTTA-3’) and the viral LTR (5’-GTGCGCGCTTCAGCAAG-3’), 1X GoTaq reaction buffer (Promega), dNTP nucleotide mix containing 200 M concentrations of each nucleotide (Promega), and 1.25U of GoTaq DNA polymerase (Promega) under the following thermocycling conditions: initial incubation at 95°C for 5 min; followed by 23 cycles at 94°C for 30s, 55°C for 30s, and 72°C for 2 min; and a final incubation at 72°C for 10 min. The second-round qPCR designed to amplify only the viral LTR-specific region contained a 1/10 volume of the first-round PCR products as the template DNA, iTaq Universal Probe Supermix (Bio-Rad), 300nM concentrations (each) of the viral LTR-specific primers that target the R region (5’-TCTGGCTAACTAGGGAACCCA-3’) and the U5 region (5’-CTGACTAAAAGGGTCTGAGG-3’), and a 100nM concentration of TaqMan probe (5’-[6-FAM]-TTAAGCCTCAATAAAGCTTGCCTTGAGTGC-[TAMRA]-3’). The qPCR run included an initial incubation at 95°C for 3 min, followed by 39 cycles of amplification and acquisition at 94°C for 15 s, 58°C for 30 s, and 72°C for 30 s. During the second round qPCR, a standard curve was generated in parallel and under the same conditions using 10-fold serial dilutions of known copy numbers (10^0^ to 10^8^) of the pNL4.3 HIV-1 molecular clone. Data were analyzed using CFX Manager software (Bio-Rad), and integrated viral DNA copy numbers were determined by plotting the qPCR data against the standard curve. To determine the specific PIC integration activity (i.e., ratio of integrated viral DNA copy numbers to corresponding PIC-associated viral DNA copy numbers) of the *in vitro* integration reactions, the PIC-associated viral DNA was isolated and copy numbers were determined by qPCR. Data were analyzed using CFX Manager software (Bio-Rad), and the viral DNA copy numbers were determined by plotting the qPCR data against the standard curve generated, as described above. All representative data are reported as means of triplicates from at least 3 independent experiments.

### Statistical analysis

All experiments were conducted at least three times with triplicates. Data were expressed as mean ± standard error of mean (SEM) obtained from three independent experiments. Significance of differences between control and treated samples were determined by Student’s *t-*test. The difference between groups were determined by paired, two-tailed student’s t-test. A p-value of < 0.05 was considered statistically significant.

## Results

### CypA depletion reduces HIV-1 DNA integration into host chromosomes

CypA regulates the early steps of reverse transcription, nuclear entry and integration site selection during HIV-1 infection. However, the direct role of CypA on post-nuclear entry steps of infection is not fully understood. Since an operationally intact HIV-1 capsid enters the nucleus and CypA specifically binds to the capsid, we hypothesized that CypA regulates the post-nuclear entry step of HIV-1 DNA integration into the host chromosomes. To test this hypothesis, we inoculated parental (CypA^+/+^) and CypA-depleted (CypA^-/-^) Jurkat cells with three different concentrations (as measured by p24/CA levels) of GFP-reporter expressing pseudotyped HIV-1 particles. 24 hours post infection, cells were harvested to measure infectivity by flow cytometry-based measurement of GFP expression. Additionally, DNA was extracted from these cells to measure levels of late reverse transcription products, 2-LTR circles as markers of nuclear entry, and proviral integration by qPCR. Results from these analyses show that CypA-depletion significantly reduced HIV-1 infectivity compared to the CypA-expressing cells (Fig. 1A). Particularly, the ∼30% of the CypA^+/+^ cells were positive for GFP, whereas only 12% of CypA^-/-^ cells expressed GFP, resulting in a >2-fold reduction in infectivity. This level of reduction was sustained despite of the amount of virus particles used (data not shown). As expected, inhibiting either reverse transcription by an RT inhibitor or an integrase inhibitor, robustly reduced infection.

**Figure 1:**
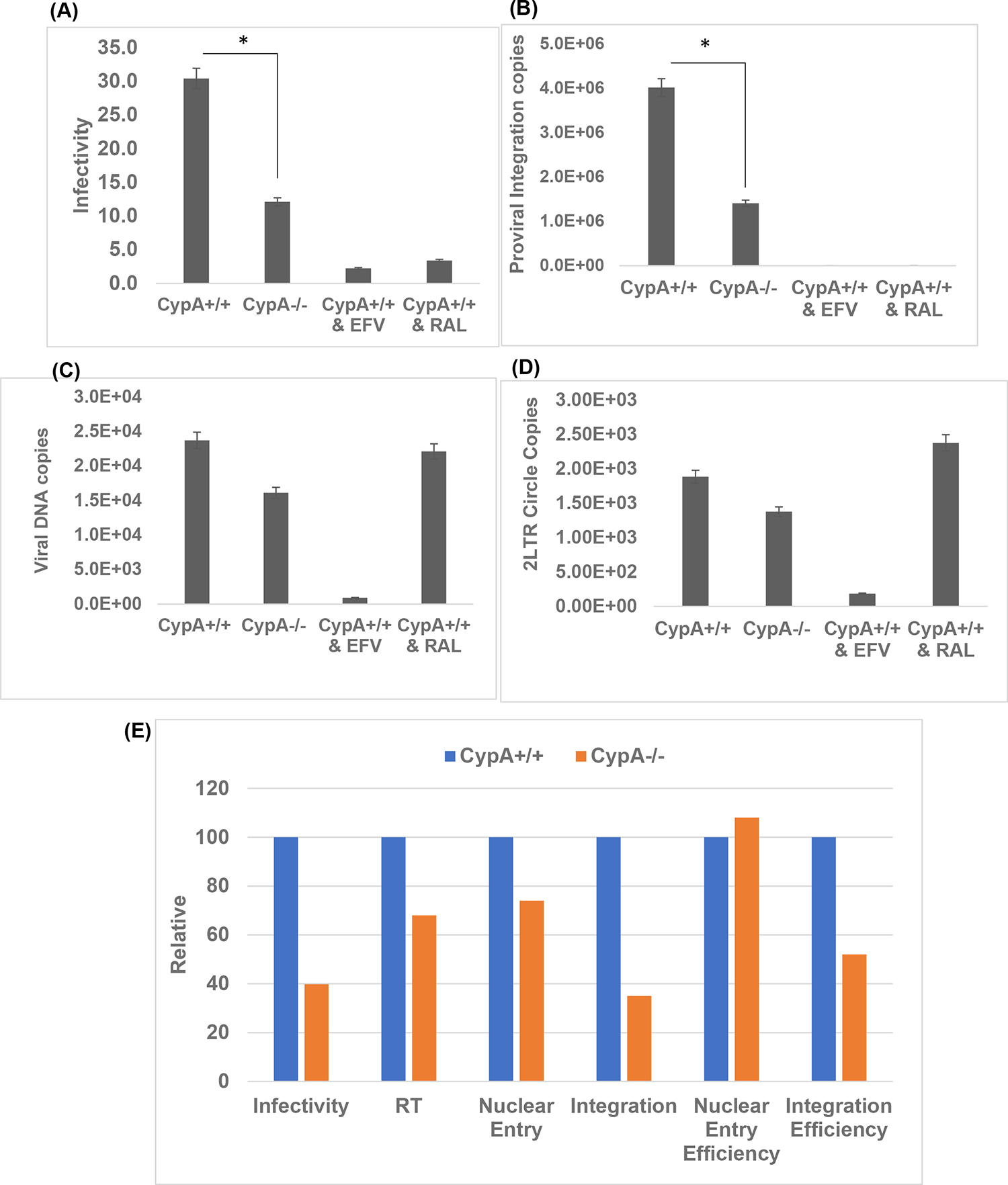
CypA-depletion reduces HIV-1 integration. **(A) Effects of CypA-depletion on HIV-1 infectivity.** Parental (CypA+/+) Jurkat T cells spinoculated with HIV-1.GFP reporter virus in the absence or presence of efavirenz and raltegravir, and the CypA-/- Jurkat T cells spinoculated with HIV-1.GFP reporter virus were all cultured for 24 hours and the intracellular GFP fluorescence (marker of infectivity) was determined by FACS. **(B) Effects of CypA-depletion on HIV-1 integration.** Integrated viral DNA (proviral DNA) were quantified by *Alu-gag* nested PCR assay of total DNA isolated from infected cells. Copy numbers were calculated from standard curves generated in parallel using 10-fold serial dilutions of known copy numbers (10^0^ - 10^8^) of a HIV-1 proviral molecular clone during the 2^nd^ round qPCR. **(C-D) Effects of CypA-depletion on HIV-1 reverse transcription and nuclear entry.** Late RT products (C) and 2-LTR circles (D) in the total DNA were quantified by qPCR. Copy numbers were calculated from standard curves generated in parallel using 10-fold serial dilutions of known copy numbers (10^0^ - 10^8^) of a HIV-1 proviral molecular clone (for late RT products) or the p2LTR plasmid (for 2-LTR circles). **(E) Summary comparison of the effects of CypA-depletion on HIV-1 reverse transcription, nuclear entry and proviral DNA integration**. To analyze specific effects of CypA on HIV-1 integration, the data of CypA-depletion on infection (Panel A), proviral integration (Panel B) reverse transcription (Panel C), and 2-LTR circles (Panel D), were plotted as percentage relative to the parental (CypA^+/+^) cells. Effect of CypA-depletion on nuclear entry efficiency was determined by calculating the ratio of copy number of 2-LTR circles to the respective late RT products, whereas proviral integration efficiency was determined by calculating the ratio of copy number of integrated viral DNAs to the corresponding late RT products. Data shown are mean values from three independent experiments with error bars representing the standard error of the mean (SEM). The p-values (*) represents statistical significance (p< 0.05).

To probe the infectivity defect in CypA^-/-^ cells, we staged the early steps of infection by qPCR using DNA from infected CypA^+/+^ and CypA^-/-^ Jurkat cells. First, we measured the copy numbers of the integrated viral DNA (proviral DNA) through a standard curve generated using the proviral molecular clone in the second round qPCR. Interestingly, we observed a significant reduction in the levels of proviral DNA integration into the chromosomes of CypA^-/-^ cells compared to the integration levels in the CypA^+/+^ cells (Fig. 1B). Notably, the level of reduction in proviral integration (∼50%) was quantitatively comparable to the decreased infectivity observed in the CypA^-/-^ cells (Fig. 1A). These observations suggested that the block at the integration step contributes to the infectivity defect in CypA^-/-^ cells. However, HIV-1 integration is dependent on viral DNA synthesis by the RTC and import of the viral DNA into the nucleus of the infected cell. Since, CypA is known to influence both viral DNA synthesis and nuclear entry, we estimated the levels of late reverse transcripts (LRT) and 2-LTR circles, as a marker of nuclear entry, in these cells by qPCR. Copies of LRT and 2-LTR circles were estimated through a standard curve generated using the proviral molecular clone and 2-LTR clone, respectively. These measurements revealed that the levels of viral DNA were reduced in CypA-depleted cells when compared to the parental cells (Fig. 1C). Similarly, the copies of 2-LTR products were also lower in CypA^-/-^ cells relative to the CypA-expressing cells (Fig. 1D). These reductions in viral DNA synthesis and 2-LTR circles are in accordance with published reports that CypA is required to promote early steps of reverse transcription and nuclear entry [32, 103]. To tease-out whether the reduced RT and nuclear entry in contributes to the block at the integration step in CypA^-/-^ cells, we calculated the efficiency of nuclear entry through the ratio of RT copies over 2-LTR copies and proviral integration by calculating the ratio of LRT copies to proviral integration copies (Fig. 1E). Interestingly, these analyses revealed that lack of CypA minimally altered the efficiency of nuclear entry of the synthesized viral DNA. However, CypA-depletion reduced the integration efficiency of the viral DNA into the host chromosomes, by an extent that was comparable to the infectivity defect. Collectively, these results suggest that CypA promotes the post-nuclear entry step of HIV-1 integration.

### Reduced HIV-1 DNA integration due to CypA depletion is not influenced by TRIM5α

In addition to promoting HIV-1 reverse transcription and nuclear entry, CypA is known to counter the antiviral effects of Tripartite motif protein isoform 5 alpha (TRIM5α) [67, 68]. TRIM5α is a cellular restriction factor and is a potent antiviral protein that restricts infection by HIV-1 and other retroviruses [104]. Particularly, TRIM5α recognizes the lattice of the retrovirus capsid in a species-specific manner and induces premature disassembly of the viral capsid [104]. Recently, it has been demonstrated that CypA prevents binding of TRIM5α to the HIV-1 capsid [67, 68]. Therefore, we tested whether the reduced HIV-1 integration in CypA-depleted cells is dependent on the presence and absence of TRIM5α. We conducted infectivity and qPCR assays using parental (CypA^+/+^/TRIM5α^+/+^), CypA-depleted (CypA^-/-^/TRIM5α^+/+^), and double knock-out (CypA^-/-^/TRIM5α^-/-^) Jurkat cells (Fig. 2). As expected, lack of CypA significantly reduced infectivity when compared to the parental cells (Fig. 2A). Interestingly, this reduction in infectivity was sustained in cells lacking both CypA and TRIM5α, strongly suggesting that CypA’s effect on infectivity is not influenced by the presence or absence of TRIM5α. Similarly, the estimation of proviral integrant copies by Alu-based nested qPCR revealed that the presence or absence of TRIM5α has minimal effect on the reduced HIV-1 integration observed with CypA-depletion (Fig. 2B). Then, measurement of the copies of LRT and 2-LTR levels in all the three cell types by qPCR (Fig. 2C-D). The lack of TRIM5α markedly increased the copies of LRT when compared to the cells without CypA (Fig 2C). Interestingly, the level of viral DNA in double knock-out cells was comparable or slightly higher than the parental cells, suggesting that viral DNA can be efficiently synthesized in cells lacking both CypA and TRIM5α. Similarly, the levels of 2-LTR copies were also increased in double KO cells when compared to the CypA-depleted cells, but was comparable to the parental cells (Fig. 2D). This is not surprising since viral DNA synthesis in the double KO cells was robust and nuclear entry is largely dependent on copies of viral DNA synthesized in an infected cell. Collectively, these results indicate that lack of TRIM5α minimally affects the detrimental effect of CypA-depletion on HIV-1 integration.

**Figure 2:**
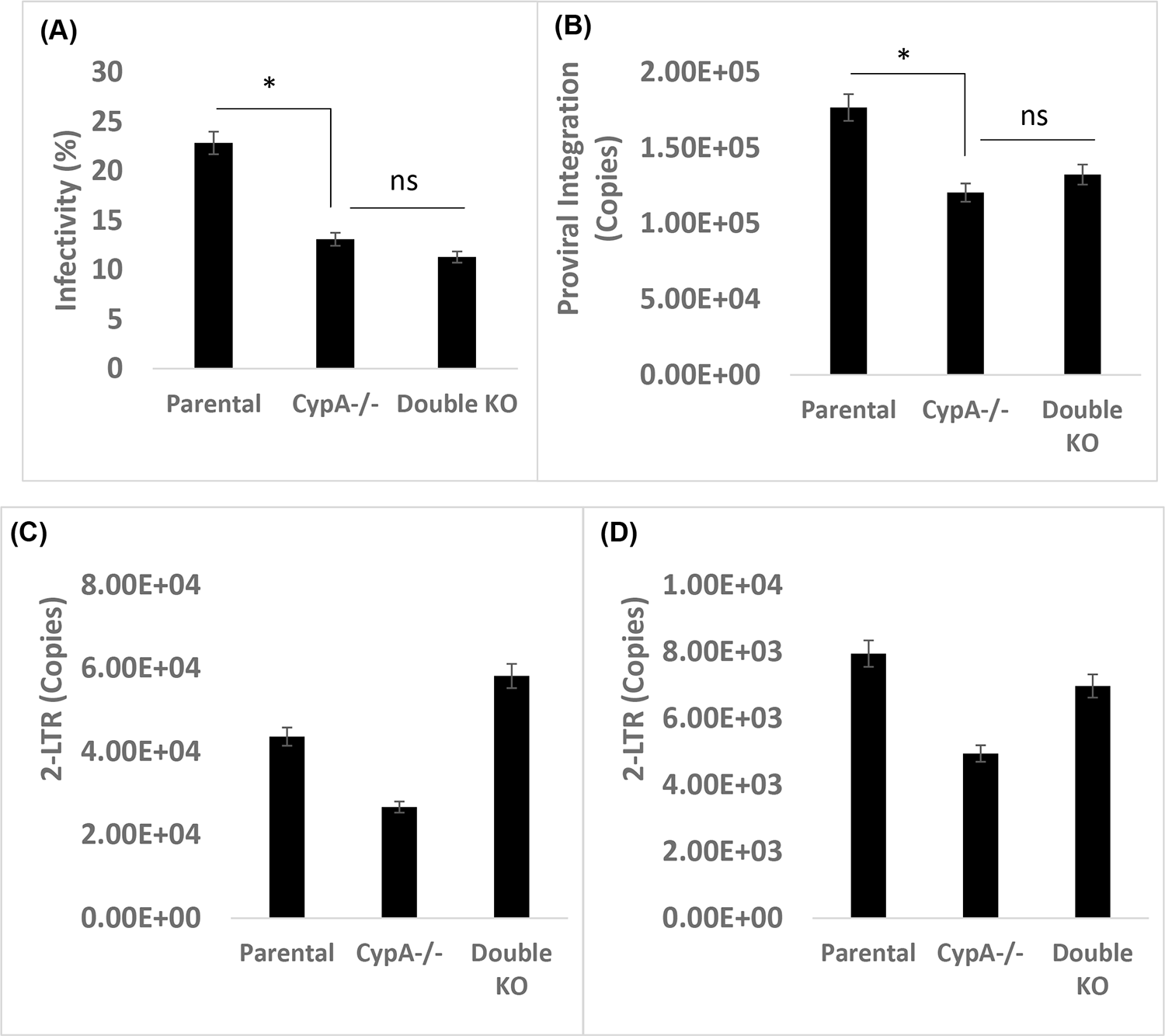
Reduced infectivity and proviral integration in CypA-depleted cells are sustained in cells lacking TRIM5α. **(A) Effects of CypA- and TRIM5α-depletion on HIV-1 infectivity.** Parental (CypA+/+), CypA-/-, and double knockout (CypA-/- & TRIM5α-/-) Jurkat T cells spinoculated with HIV-1.GFP reporter particles were cultured for 24 hours and the intracellular GFP fluorescence (marker of infectivity) was determined by FACS. **(B) Effects of CypA- and TRIM5α-depletion on HIV-1 integration.** Proviral DNA were quantified by *Alu-gag* nested PCR assay of total DNA isolated from infected cells and copy numbers were calculated from standard curves generated in parallel during the 2^nd^ round qPCR. **(C-D) Effects of CypA- and TRIM5α-depletion on HIV-1 reverse transcription and nuclear entry.** Late RT products (C) and 2-LTR circles (D) in the total DNA were quantified by qPCR and copy numbers were calculated from appropriate standard curves generated in parallel. Data shown are mean values from three independent experiments with error bars representing the SEM. The p-values (*) represents statistical significance (p< 0.05) and ns stands for nonsignificant.

### Inhibition of CypA-capsid binding by Cyclosporin A blocks HIV-1 integration

Our results in Figs. 1-2 demonstrated that lack of CypA blocks HIV-1 integration independent of reverse transcription and nuclear entry. To further probe CypA’s role in HIV-1 integration, we next used a pharmacological inhibitor cyclosporin A (CsA) that is known to inhibit CypA-CA binding. CsA is a cyclic undecapeptide [105, 106] that inhibits the PPIase activity of CypA by binding to the substrate-binding site of CypA [55]. It is well-established that CsA inhibits HIV-1 infection in a CypA-dependent manner [62, 65], thus serves as an appropriate probe to study CypA’s role during infection. First, we measured the antiviral activity of CsA in CypA^+/+^ and CypA^-/-^ Jurkat cells (Fig. 3A). We observed significant reduction in (∼2-fold) in percent of cells expressing GFP in CypA-depleted cells compared to the parental cells. In contrast, CsA treatment showed no effect on GFP expression in CypA-depleted cells, confirming that the antiviral activity of CsA is dependent on the presence of CypA in the infected cells. Notably, estimation of proviral integrants in these cells showed that CsA treatment significantly lowered the copies of HIV-1 DNA integrated into the host chromosomes in the CypA-expressing cells (Fig. 3B). Interestingly, in the CypA-depleted cells similar levels of integration was detected in both untreated and CsA-treated cells. As expected, the copies of integrants in CypA-depleted cells were lower compared to the untreated CypA-expressing cells. Then, to test whether the reduced integration in CsA-treated cells is dependent on lower RT and nuclear entry of the virus, we measured copies of LRT and 2-LTR circles, respectively (Fig. 3C-D). As expected, CsA reduced the number of viral DNA copies in CypA-expressing cells and had no such effect in CypA-lacking cells (Fig. 3C). Additionally, the copies 2-LTR were also lower only in CsA-treated CypA-expressing cells but not in the CypA-depleted cells (Fig. 3D). Nevertheless, the ratio between LRT copies to 2-LTR copies indicated that CsA minimally reduces in the efficiency of nuclear entry of the viral DNA (Fig. 3E). However, integration efficiency was markedly reduced upon CsA treatment in CypA-expressing cells (Fig. 3E), an effect that mirrored the inhibitory effect of this drug on infectivity. These results suggested that inhibiting CypA had a detrimental effect on HIV-1 integration and further supporting CypA’s role in post-nuclear entry step.

**Figure 3.**
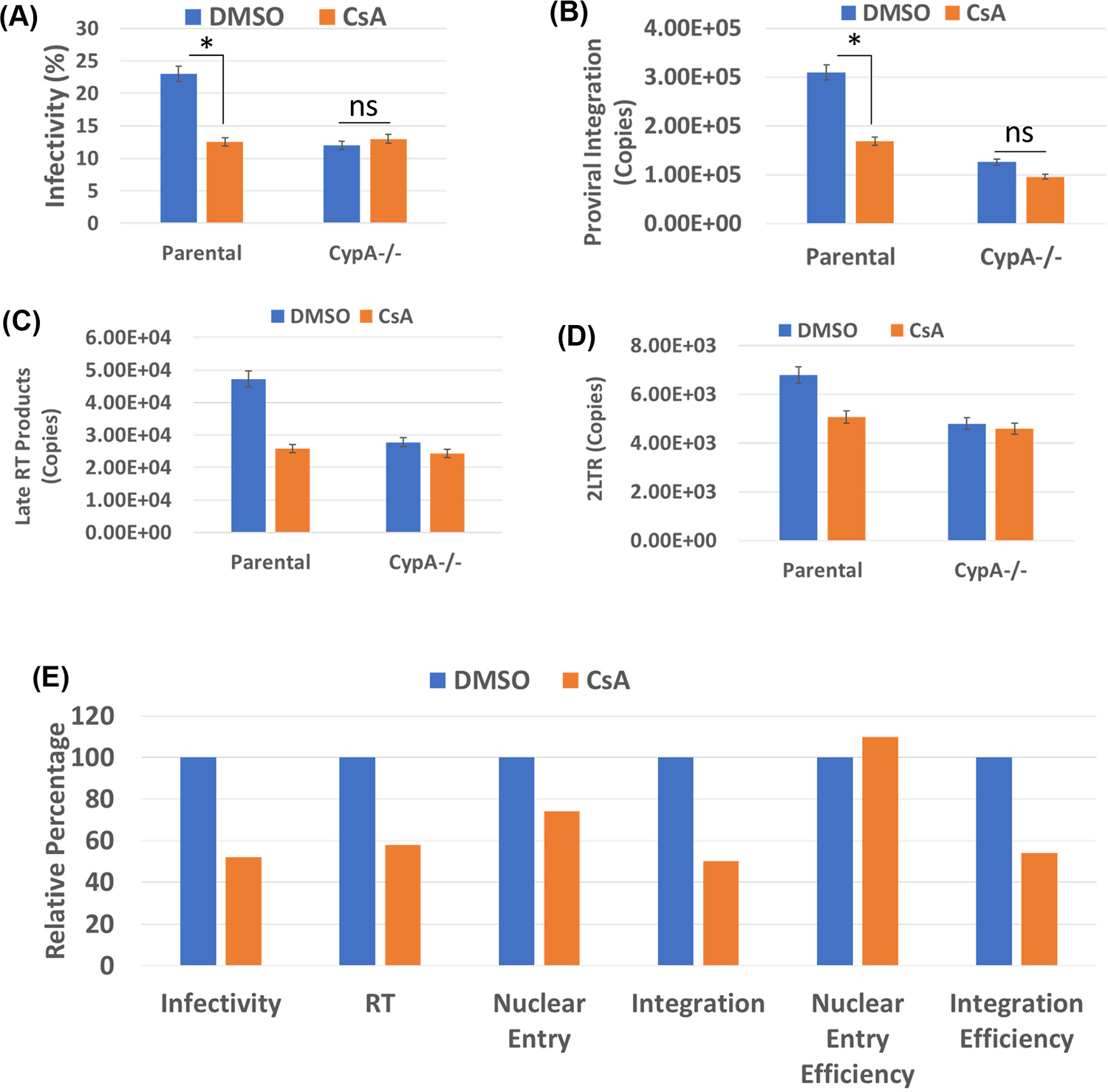
CsA markedly reduces HIV-1 integration. (A) Effects of CsA on HIV-1 infectivity. Parental (CypA^+/+^) and CypA-depleted (CypA^-/-^) Jurkat T cells inoculated with HIV-1.GFP particles in the presence of DMSO or 10 µM CsA were cultured for 24 hours and the intracellular GFP fluorescence (indicator of infectivity) was determined by FACS. **(B) Effects of CsA on HIV-1 integration.** Integrated viral DNA were quantified by *Alu-gag* nested PCR assay of total DNA isolated from infected cells and copy numbers were calculated from standard curves generated in parallel during the 2^nd^ round qPCR. **(C-D) Effects of CsA on HIV-1 reverse transcription and nuclear entry.** Late RT products (C) and 2-LTR circles (D) in the total DNA from infected cells were determined by qPCR and copy numbers were calculated from standard curves generated in parallel. **(E) Summary comparison of the effects of CsA on early steps of HIV-1 infection**. To analyze specific effects of CsA on HIV-1 integration, the data of CsA’s effect on virus infectivity (Panel A), proviral integration (Panel B) reverse transcription (Panel C), and nuclear entry (Panel D), were plotted as percentage relative to that of DMSO-treated cells. Nuclear entry efficiency and proviral integration efficiency were calculated as described in Fig. 1 legend. Data are mean values from at least three independent experiments with error bars representing the SEM. The p-values (*) represents statistical significance (p< 0.05).

### HIV-1 CA mutants that fail to bind to CypA are blocked at integration step

Our molecular and pharmacologic studies demonstrated that CypA is required for optimal HIV-1 integration (Figs. 1-3). To further assess CypA’s function at this post-nuclear entry step of HIV-1 infection, we employed HIV-1 CA mutants P90A and G89V in our study. These two mutations are located in the CypA binding loop of HIV-1 CA and capsids of these mutant viruses fail to bind to CypA [107]. Nevertheless, these mutants retain the ability to infect target cells albeit at lower level [67, 68] making these CA mutants useful tools to study CypA’s role during HIV-1 infection. We inoculated CypA^+/+^ and CypA^-/-^ Jurkat cells with HIV-1 G89V (Fig. 4A) or P90A particles (Fig. 4B). A significant reduction (∼2-fold) in infectivity was observed with each of these mutants when compared to the wild-type CA containing particles in CypA-expressing cells, consistent with published studies [67, 68]. In contrast, in CypA-depleted cells the infectivity levels of the mutant viruses were comparable to the wild type virus, confirming that the infectivity defect of these mutants is mapped to CypA. As expected, HIV-1 particles with wild-type CA showed a 2-fold reduced infectivity in CypA-depleted cells compared to the CypA-expressing cells (Fig. 4A-B).

**Figure 4.**
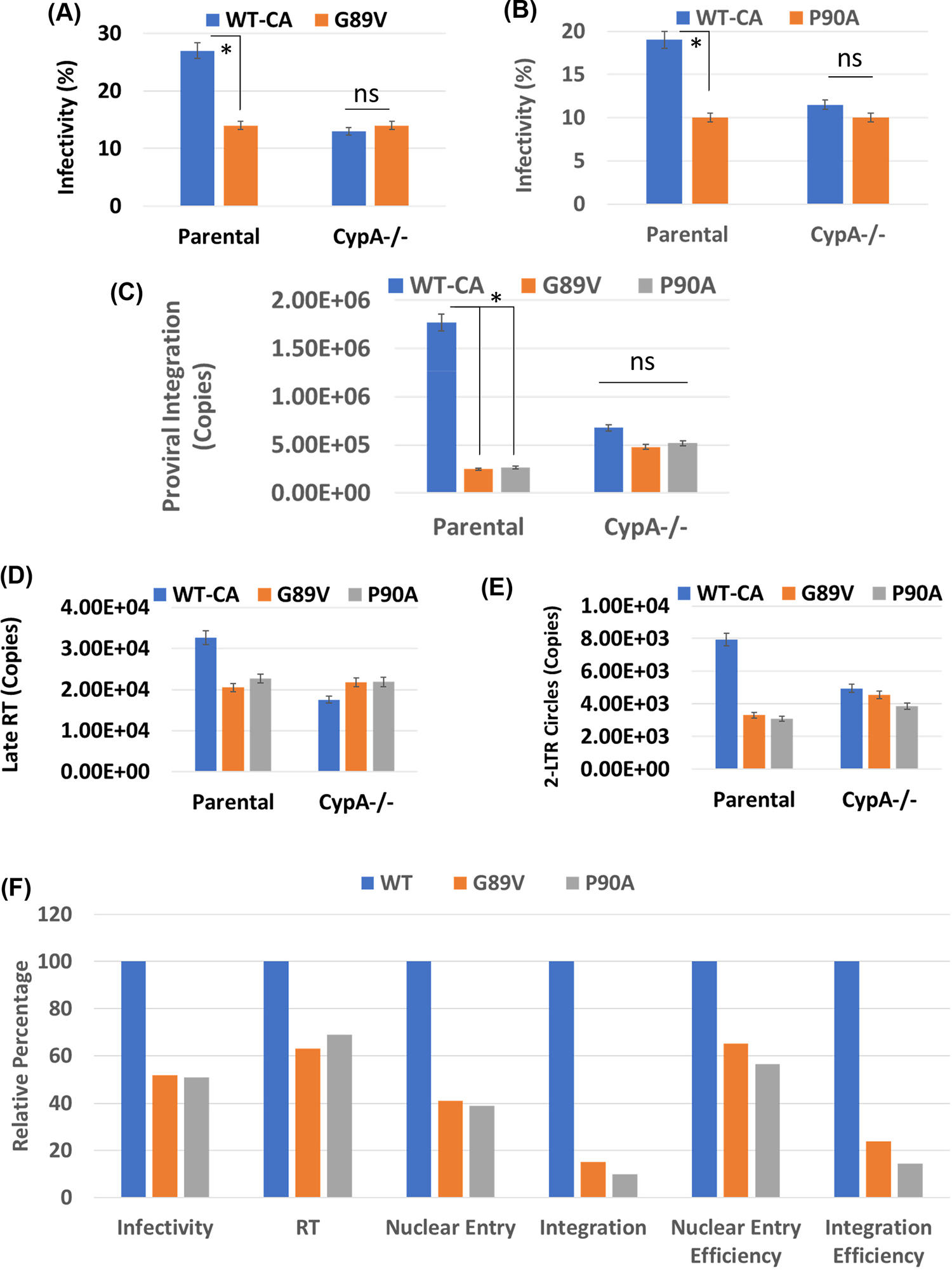
HIV-1 CA mutant viruses with disrupted CypA binding are defective in proviral DNA integration. Effects of CA mutations (A) G89V and (B) P90A on HIV-1 infectivity. Parental (CypA^+/+^) and CypA-depleted (CypA^-/-^) Jurkat T cells inoculated with wild-type CA or mutant CA HIV-1 particles were cultured for 24 hours and intracellular GFP fluorescence (indicator of infectivity) was determined by FACS. **(C) Effects of CA mutations G89V and P90A on HIV-1 integration.** Integrated viral DNA were quantified by *Alu-gag* nested PCR assay of total DNA from infected cells and copy numbers were calculated from standard curves generated in parallel during the 2^nd^ round qPCR. **(D-E) Effects of CA mutations G89V and P90A on HIV-1 reverse transcription and nuclear entry.** Late RT products (D) and 2-LTR circles (D) in the total DNA from infected cells were quantified by qPCR and copy numbers were calculated from standard curves generated in parallel. **(F) Summary comparison of the effects of CA mutations with disrupted CypA binding on early steps of HIV-1 infection.** To analyze specific effects of G89V or P90A on HIV-1 integration, the data of CA mutations’ effect on virus infectivity (Panel A-B), proviral integration (Panel C) reverse transcription (Panel D), and nuclear entry (Panel E), were plotted as percentage relative to that of WT CA HIV-1 control. Nuclear entry efficiency and proviral integration efficiency were calculated as described in Fig. 1. Data are mean values from at least three independent experiments with error bars representing the SEM. The p-values (*) represents statistical significance (p< 0.05).

Then, we examined the early steps at which the infection of G89V and P90A viruses are impaired by measuring proviral integration, LRT, and 2-LTR copies (Fig. 4C-E). Notably, the proviral integration levels of each of these mutant viruses were significantly lower when compared to the wild-type CA virus in CypA-expressing cells (Fig. 4C). As expected, in CypA-depleted cells the mutant viruses and the wild-type CA virus integrated at a comparable level, although the copies of integrated wild-type HIV-1 CA DNA were lower compared to the numbers in CypA-expressing cells. To study whether the reduced integration of the mutant viruses is dependent on lower RT and nuclear entry, we measured copies of LRT and 2-LTR, respectively, in CypA^+/+^ and CypA^-/-^ Jurkat cells (Fig. 4D-E). Viral DNA synthesis in cells inoculated with either G89V or P90A viruses was lower in CypA-expressing cells when compared to the wild-type CA virus (Fig. 4D). Yet, in CypA-depleted cells the levels of viral DNA of the mutant viruses are comparable to the wild-type virus (Fig. 4D). In addition, the copies 2-LTR circles formed in the nucleus of the cells infected with either of the mutant virus was lower compared to the wild-type CA virus (Fig. 4E). Interestingly, the efficiency of nuclear entry of the mutant viruses was also reduced in CypA-expressing cells (Fig. 4F). Intriguingly, the efficiency of integration of both G89V and P90A was also markedly reduced in CypA-expressing cells, an effect that is quantitatively lower than the nuclear entry efficiency (Fig. 4F). Collectively, these results suggest that HIV-1 CA mutants that lack the ability to engage with CypA are blocked at nuclear entry and HIV-1 integration, further supporting that binding of CypA to the viral capsid in target cells promotes HIV-1 integration.

### HIV-1 PICs from CypA-depleted cells retain lower viral DNA integration activity

Integration of HIV-1 DNA into the host chromosomes in an infected cell is carried out by the PIC-the biochemically machinery that consists of the viral DNA, integrase and other host/viral factors [24]. Importantly, PICs retain integration activity *in vitro* after extraction from acutely infected cells and have been extensively used to study mechanism of retroviral integration [94, 108, 109]. Therefore, to determine whether lower integration in CypA-depleted cells is a consequence of altered PIC function, we prepared cytoplasmic PICs from Jurkat cells inoculated with DNase I-treated HIV-1 particles [93, 94, 108–110]. In parallel, we prepared cytoplasmic extracts from uninfected cells as negative controls. These extracts were were assayed for integration *in vitro* into a 2 kb linear target DNA and the integrants were quantified by nested qPCR [93, 94, 108–110]. The results in Fig. 5A show that the extracted PICs from the parental CypA-expressing cells productively carry out integration of the viral DNA into the target DNA. Notably, the PIC integration activity was significantly inhibited in the presence of RAL, an integrase inhibitor, demonstrating specificity of the integration assay. Furthermore, the extracts from the uninfected cells and the lack of PICs or target DNA in the reaction mixture exhibited no measurable integration activity, further establishing the specificity of the PIC-associated viral DNA integration activity measurements.

**Figure 5.**
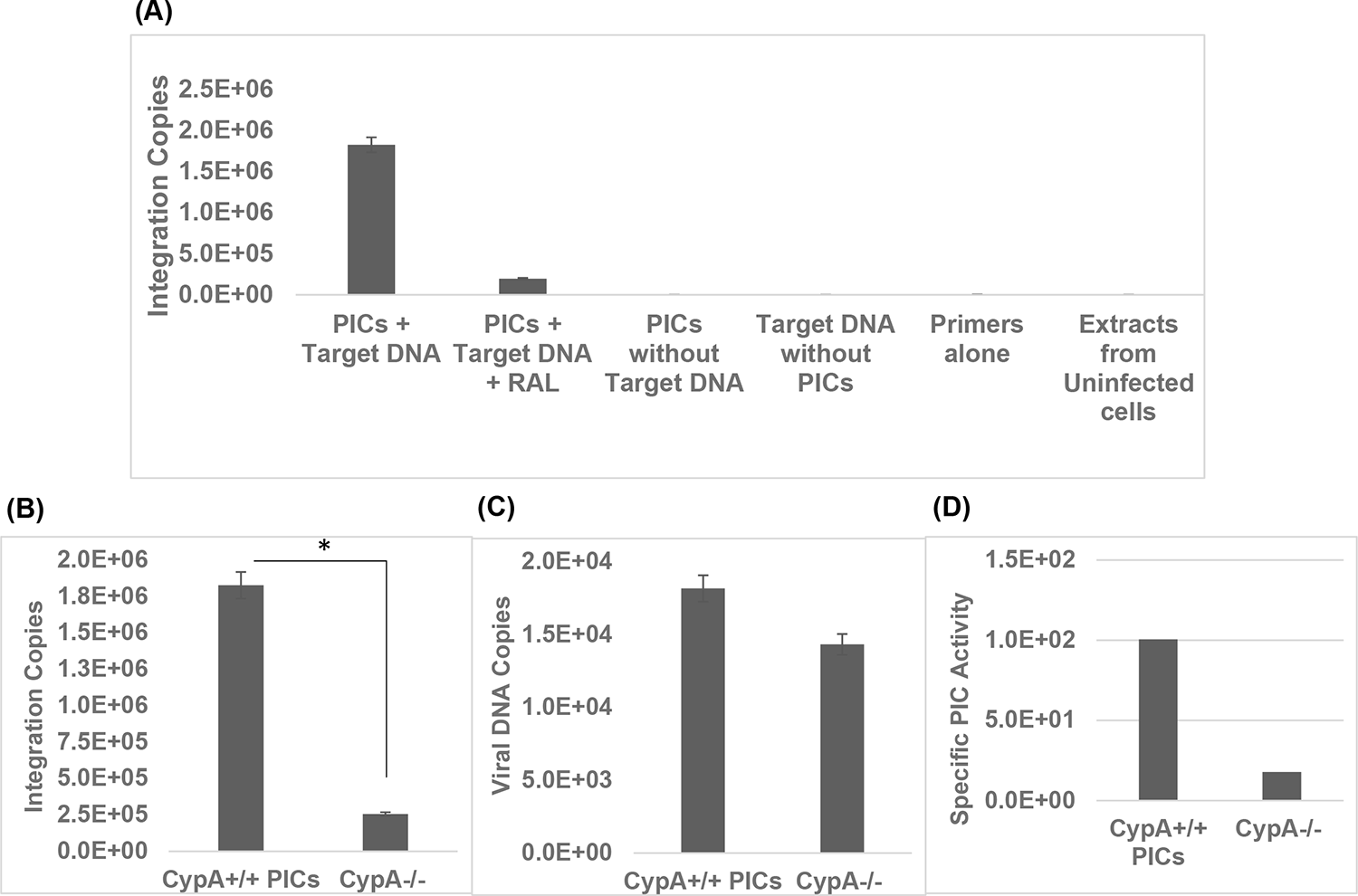
Effect of CypA-depletion on the integration activity of HIV-1 PICs. (A) PIC-associated integration activity measurements in vitro. **(A)** Parental (CypA^+/+^) Jurkat T cells spinoculated with high titer HIV-1 particles (1500 ng/mL of p24) were cultured for 5 hours followed by extraction of PIC-containing cytoplasmic extracts (Cy-PICs). In vitro integration assays using the Cy-PICs as the source of integration activity and quantification of viral DNA integration by nested PCR were carried out with appropriate controls. **(B) Integration activity of PICs.** In vitro integration assays of Cy-PICs extracted from CypA-expressing (CypA^+/+^) parental cells and from CypA-depleted (CypA^-/-^) Jurkat T cells were carried out as described in materials and methods and the copy numbers of integrated viral DNAs were determined by nested PCR that included generation of standard curves in parallel during the 2^nd^ round qPCR. **(C) Viral DNA content of the PICs.** Copy numbers of viral DNA in the extracted PICs were quantified by qPCR and copy numbers were calculated from standard curves generated in parallel. **(D) Specific PIC activity.** The specific PIC activity was determined by calculating the ratio of integrated viral DNA copy numbers (B) to the corresponding PIC-associated viral DNA copy numbers (C). Mean values from three independent experiments are shown with error bars representing the SEM. The p-values (*) represents statistical significance (p< 0.05).

Next, we analyzed the effects of CypA on PIC-associated viral DNA integration. First, PICs were extracted from CypA^+/+^ and CypA^-/-^ Jurkat cells inoculated with HIV-1 particles. The activity of these Cy-PICs was assayed *in vitro*, and viral DNA integration was quantified as described in Fig. 1A. Surprisingly, the Cy-PICs from (CypA^-/-^) cells displayed significantly lower integration activity relative to the Cy-PICs from CypA^+/+^ control cells (Fig. 5B). To clarify whether the lower integration activity is due to reduced number of PICs formed in these cells, we quantified the amount of viral DNA in the Cy-PICs from both the cells types. Interestingly, the level of viral DNA in CypA^-/-^cells was slightly lower (Fig. 5C). However, calculation of specific PIC activity considering the ratio of integration copies over viral DNA copies, revealed that integration activity of the PICs was significantly reduced in CypA^-/-^ cells compared to the control cells (Fig. 5D). These observations suggested that lack of CypA reduces viral DNA integration by the PIC and this reduction is not due to reduced number of PICs formed in these cells.

### Lower integration activity of HIV-1 PICs from CypA-depleted cells are not affected by the presence or absence of TRIM5α

CypA has been shown to counter the antiviral function of TRIM5α by preventing its binding to the HIV-1 capsid [67, 68]. Given our observation that CypA-depletion reduces PIC-integration activity (Fig. 5), we next probed whether this phenotype is affected by the presence and absence of TRIM5α. We extracted PICs from acutely infected parental (CypA^+/+^/TRIM5α^+/+^), CypA-depleted (CypA^-/-^/TRIM5α^+/+^), and double knock-out (CypA^-/-^ /TRIM5α^-/-^) Jurkat cells (Fig. 6). In accordance with our results in Fig. 5, PICs recovered from cells lacking CypA retained significantly lower integration activity relative to the PICs from parental cells expressing both CypA and TRIM5α (Fig. 6A). However, PICs extracted from cells lacking both CypA and TRIM5α also retained lower integration activity compared to the parental cells, but showed similar levels of activity relative to the CypA-depleted cells. Then, measurement of the copies of viral DNA in these PICs indicated that in cells lacking TRIM5α the copies of LRT are comparable to the parental cells but are higher relative to CypA-depleted cells (Fig 6B). Nevertheless, calculation of PIC-specific activity indicated that lack of TRIM5α minimally alters the activity of PICs compared to those from cells lacking CypA (Fig. 6C). Thus, the effects of CypA on PIC-integration activity appears to be unrelated to TRIM5α.

**Figure 6:**
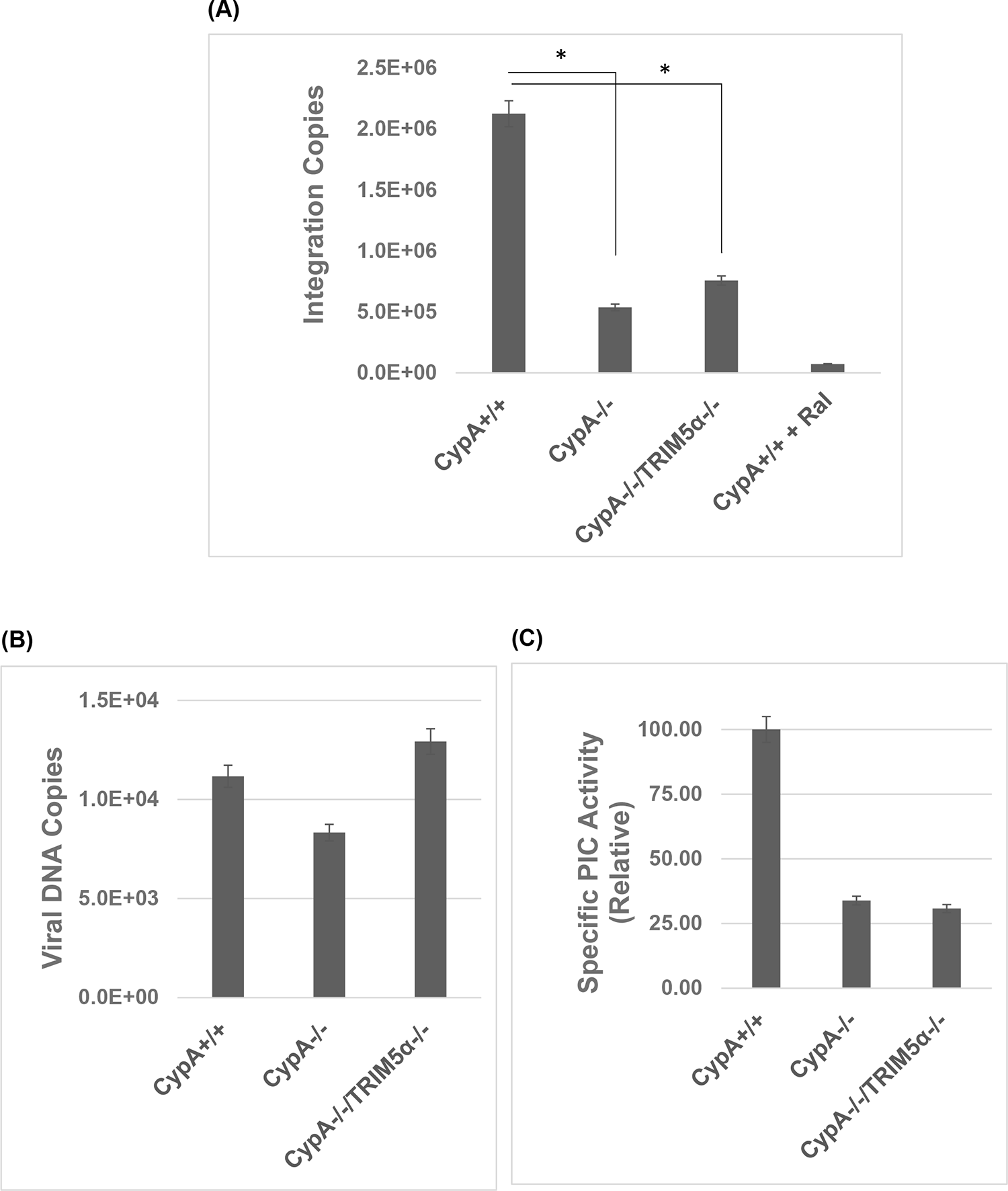
CyA depletion-associated reduction in PIC integration activity is sustained in PICs from cells lacking TRIM5α as well. **(A) Integration activity of PICs.** Parental (CypA+/+), CypA-/-, and double knockout (CypA-/- & TRIM5α-/-) Jurkat T cells spinoculated with HIV-1 particles were cultured for 5 hours followed by extraction of Cy-PICs. In vitro integration assays of Cy-PICs were carried out as described in materials and methods and the copy numbers of integrated viral DNAs were determined by nested PCR that included generation of standard curves in parallel during the 2^nd^ round qPCR. The copy numbers of integrated viral DNAs were determined by nested PCR that included generation of standard curves in parallel during the 2^nd^ round qPCR. **(B) Viral DNA content of the PICs.** Copy numbers of viral DNA in the extracted PICs were quantified by qPCR and copy numbers were calculated from standard curves generated in parallel. **(C) Specific PIC activity.** The specific PIC activity was determined by calculating the ratio of integrated viral DNA copy numbers (B) to the corresponding PIC-associated viral DNA copy numbers (C) and plotted as percentage relative to specific activity of Parental (CypA+/+) PICs. Data shown are mean values from three independent experiments with error bars representing the SEM. The p-values (*) represents statistical significance (p< 0.05) and ns stands for nonsignificant.

### Addition of CypA protein stimulates the integration activity of HIV-1 PICs

Biochemical analysis of PICs from CypA-depleted cells established that lack of CypA reduces the integration activity of these replication complexes. To test whether the CypA directly modulates PIC function, we measured effects of recombinant CypA on PIC-integration activity. A range of concentrations of CypA (0 to 20 μM) were added to integration reaction mixtures containing PICs extracted from acutely infected parental (CypA-expressing) and CypA-depleted cells. Quantification of viral DNA integration activity illustrate that exogenous addition of CypA protein markedly stimulated integration activity of PICs extracted from both CypA-expressing and CypA-lacking cells (Fig. 7A). Particularly, addition of CypA protein up to 10 μM enhances integration activity of PICs from CypA-expressing cells in a dose dependent manner, followed by a saturating effect with the addition of 20 μM protein. In contrast, PICs extracted from CypA-depleted cells showed a dose-dependent increase with 20 μM protein (Fig. 7B). Notably, calculation of fold-increase in integration activity revealed that addition of CypA protein up to 10 μM concentration increased the activity of both PICs at a comparable level. However, addition of 20 μM CypA significantly stimulated the activity of the PICs extracted from the CypA-depleted cells when compared to those from CypA-expressing cells (Fig. 7C). These biochemical studies provide strong evidence that presence of CypA is required for the optimal integration activity of PICs.

**Figure 7.**
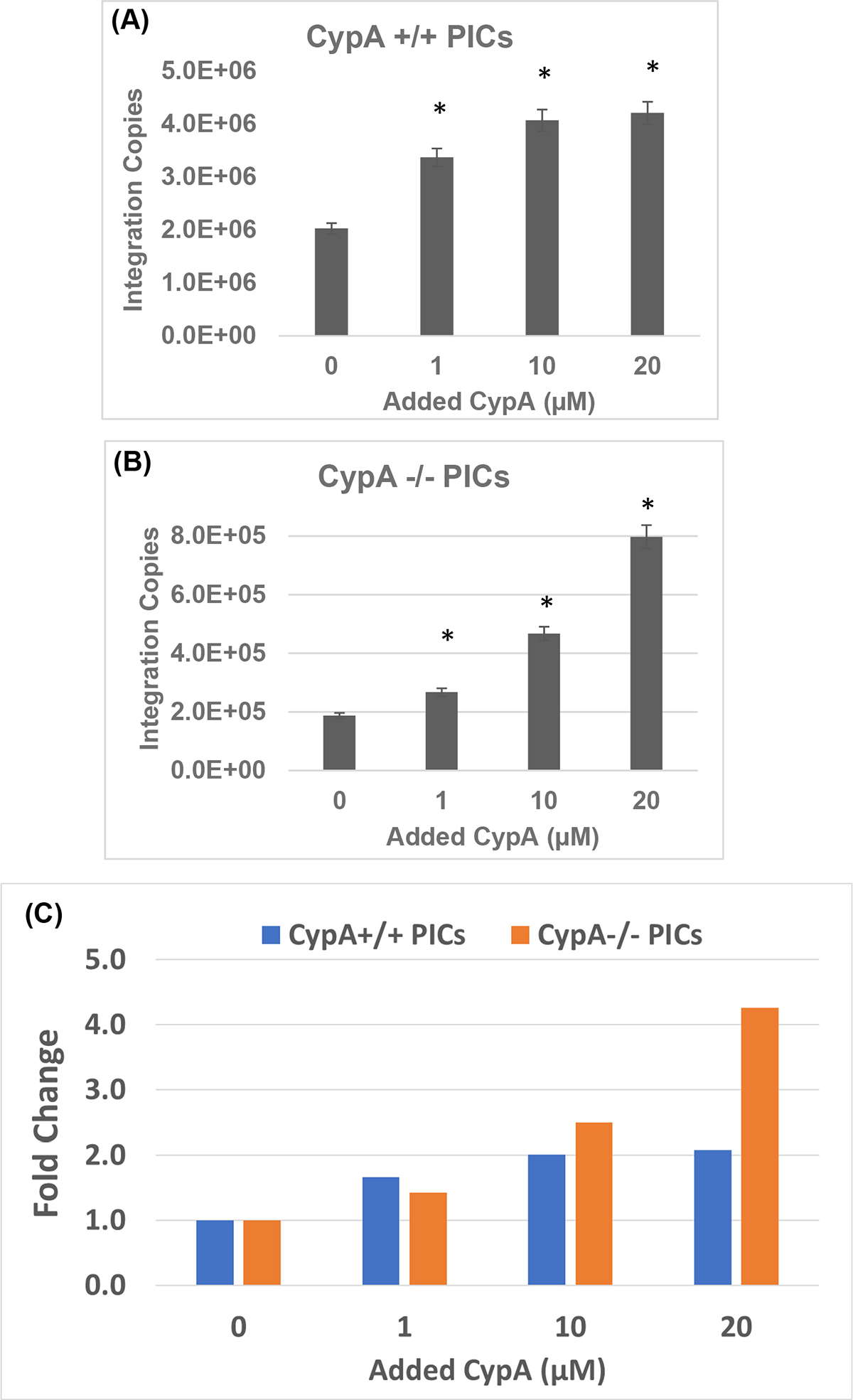
Effects of recombinant CypA protein on PIC-associated integration activity in vitro. (A-B) Integration activity measurement in the presence of recombinant CypA protein. Integration activity in vitro of PICs extracted from CypA-expressing (CypA^+/+^) parental cells **(A)** and from CypA-depleted (CypA^-/-^) Jurkat T cells **(B)** were assessed in the absence (Control) or presence of varying concentrations (0-20 µM) of purified recombinant CypA protein, and the copy number of the integrated viral DNAs were determined by nested PCR. **(C)** Integration activity plotted as fold change relative to the integration activity of the respective control PICs. Mean values from three independent experiments are shown with error bars representing the SEM. The p-values (*) represents statistical significance (p<0.05) and ns stands for nonsignificant.

## Discussion

Over forty years ago, CypA was identified as the first host factor to bind to the HIV-1 Gag polyprotein [60]. Specifically, CypA was demonstrated to bind to the proline-rich CypA-binding loop of the HIV-1 CA-NTD region [61–63]. Now, it is well-established that CypA-CA binding promotes the early steps of reverse transcription and nuclear entry during HIV-1 infection [29, 32, 61–63]. Additionally, CypA-CA binding is also required to counter the anti-viral effects of the TRIM5α protein [67, 68]. However, very little is known whether CypA-CA binding directly influences post-nuclear entry steps of HIV-1 infection. Since human CypA is predominantly a cytosolic protein, it was logical to probe a direct or functional role of CypA in HIV-1 reverse transcription and nuclear entry, steps classically known to be completed in the cytoplasmic side of an infected cell. Nevertheless, there is evidence that CypA localizes to the nucleus of *Saccharomyces cerevisiae* [111] and in human monocyte-derived macrophages [83]. CypA has also been reported to localize to the nucleus of Jurkat T-cells during cytokinesis [88] and upon stimulation of cells with stressors [89]. Most importantly, strong evidence is accumulating that an intact or near-intact HIV-1 capsid enters the nucleus of an infected cell and nuclear capsid uncoating occurs just prior to- and near the site of viral DNA integration [90–92]. Additionally, completion of HIV-1 reverse transcription is also emerging as a nuclear event. On the basis of these significant new developments, we hypothesized that CypA-CA interaction plays a direct role in post-nuclear entry steps of HIV-1 infection. Particularly, we predicted that CypA could influence HIV-1 DNA integration step by remaining engaged with the nuclear HIV-1 capsid. Indeed, our results obtained through a combination of molecular, genetic, pharmacologic, and biochemical approaches show that CypA promotes HIV-1 DNA integration into the host chromosomes. Particularly, our study demonstrates that CypA facilitates HIV-1 PIC-mediated viral DNA integration, a previously unknown function of this host factor. We also describe that CypA’s effect on HIV-1 DNA integration is independent of its role in reverse transcription and nuclear entry as well as the presence or absence of TRIM5α. Finally, our results also suggest that CypA’s effect on HIV-1 integration is dependent on its ability to bind to the viral capsid.

There is indirect evidence that the CypA-CA interaction influences post-nuclear steps of HIV-1 infection. For example, specific CA mutations (G89V or P90A) in the CypA-binding loop that disrupt CypA-CA interaction alter integration targeting into specific regions of the host genome [32]. However, the effects of these CA mutations on the levels of HIV-1 DNA integration into the host chromosomes have not been systematically measured. Recently, our group reported that CypA expression directly correlates with the levels of integration of HIV-1 CA mutant (R264K) that evades the CTLs [93]. Specifically, we showed that reduced infectivity of the R264K mutant is caused by the block at the integration step and was not a consequence of any defect at the reverse transcription or the nuclear entry step. Especially, several fold increase in proviral integration of the R264K mutant was measured in CypA-depleted cells when compared to CypA-expressing cells. These observations were surprising, since the R264K mutation is not located within or near the CypA-binding loop of HIV-1 CA. Nevertheless, our results strongly supported a functional role of CypA in post-nuclear entry steps of HIV-1 infection, particularly in the integration step of this CTL escape CA mutants. The goal of the present study was to better understand CypA’s role in the post-nuclear entry steps of infection. Therefore, we probed a direct role of CypA in HIV-1 integration using a multipronged approach of CypA-depletion (Fig. 1), use of CsA-a CypA inhibitor (Fig. 3), and CA mutants G89V and P90A (Fig. 4). Interestingly, HIV-1 DNA integration into the host chromosomes was significantly lower in cells lacking CypA when compared to cells-expressing endogenous levels of CypA (Fig. 1B). Notably, the level of reduction in proviral integration was comparable to the infectivity block in the CypA-depleted cells (Fig. 1A-B). Accordingly, by disrupting CypA-CA interaction by CsA treatment (Fig. 3) or through specific CA mutations (G89V or P90A) (Fig. 4), we also measured lower HIV-1 integration levels. Most importantly, the integration defect caused by CsA-treatment (Fig. 3B) or CA mutations (Fig. 4C) was not observed in cells lacking CypA (Fig. 3B and Fig. 4C). Collectively, these observations provide strong evidence that CypA expression is directly linked to HIV-1 DNA integration into the host chromosomes-a nuclear step of infection.

HIV-1 integration is dependent on the early step of viral DNA synthesis. Importantly, binding of CypA to the viral capsid is known to promote HIV-1 reverse transcription [29, 67, 68]. Therefore, we probed whether CypA’s effect on HIV-1 DNA integration is a consequence of an effect of this host factor on viral DNA synthesis. Accordingly, we quantified viral DNA copies in CypA-depleted cells (Fig. 1C), CsA-treated cells (Fig. 3C) and by CA mutants (Fig. 4D). As expected, viral DNA synthesis was reduced, albeit to varied levels depending on the experimental condition. Nevertheless, the overall effects were in accordance with published studies. For instance, CA specific mutations P90A or G89V/A significantly reduce viral DNA synthesis both in transformed cells [65, 66] and in human primary peripheral blood mononuclear cells (PBMCs) [69]. Notably, two recent studies describe CypA’s positive effects on HIV-1 reverse transcription in human primary macrophages and CD4+ T cells [67, 68]. For example, viral DNA synthesis by P90A was significantly reduced in primary macrophages and CD4+ T cells [67]. Similarly, reverse transcription of P90A and G89V mutants was dramatically reduced in primary CD4+ T cells [68]. The effects of CypA on HIV-1 reverse transcription have been linked to its ability to stabilize the viral capsid [70–76], even though the exact mechanism is not fully understood. Additionally, whether reduced viral DNA synthesis directly manifests similar blocks in the subsequent steps of HIV-1 infection is unclear. This is mainly because genetic, pharmacologic and microscopy studies suggest that CypA also regulates the import, docking and entry of HIV-1 replication complex through the NPC into the nucleus of an infected cell [32, 39, 77–83]. Accordingly, our 2-LTR circle measurements revealed that lack of CypA (Fig. 1D) and disrupting CypA-CA binding by either CsA (Fig. 3D) or by specific CA mutations (Fig. 4E) also reduce HIV-1 nuclear entry. The reduced nuclear entry was CypA-specific, since no such reduction in 2-LTR copies was detected in CypA-depleted cells. Interestingly, the reduction in nuclear entry was accompanied by a mostly comparable level of reduction in copies of viral DNA in the respective conditions (Figs. 1C, 3C, and 4D). Given a lack of synergistic effect, our experimental approaches could not parse out whether the nuclear entry defect by CypA-depletion or disruption of CypA-CA binding is an independent effect or is linked to the reduced levels of viral DNA. Therefore, we quantified nuclear entry efficiency based on the ratio of the copies of viral DNA to the copies of 2-LTR circles. This method provides a better measure of the effects of reduction of viral DNA on nuclear entry, when a host factor affects both the steps. These quantitative analyses clearly illustrated that nuclear entry efficiency is minimally altered in CypA-depleted cells (Fig. 1E) and CsA-treated cells (Fig. 3E). In contrast, there was a measurable reduction in nuclear entry efficiency of the CA mutant G89V and P90A particles (Fig. 4F), suggesting that a block at nuclear entry could contribute to the infectivity defect of these variants. The nuclear entry block induced by these two CA mutations is consistent with a recent report demonstrating similar nuclear entry defects in these viruses as measured by the levels of 2-LTR circles at 36 hpi in primary CD4+ T cells [68]. However, this study also reported dramatic reduction in viral DNA synthesis, thus it was unclear whether the nuclear entry defect was caused by the markedly lower viral DNA levels. Collectively, our studies indicate that the infectivity defect in CypA-depleted cells and CsA-treated cells is likely due to a block after the nuclear entry step, whereas the CA mutants are partly blocked at the nuclear entry step.

HIV-1 DNA integration in target cells is dependent on nuclear entry. Therefore, to measure a direct role of CypA on post-nuclear entry steps, we calculated the efficiency of proviral integration through the ratio of the viral DNA copies to the estimated proviral integration copies. This quantitative analysis was essential to reduce the confounding effects of reduced nuclear entry on proviral integration, particularly for the CA mutants that showed a measurable reduction in nuclear entry efficiency. These analyses illustrated that in CypA-depleted cells the efficiency of integration was markedly reduced to a level that is comparable to the infectivity defect (Fig. 1E). Given that CypA-depletion minimally reduced nuclear entry efficiency (Fig. 1E), these observations provided strong evidence for a functional link between CypA expression and HIV-1 integration. Similarly, CsA treatment also markedly reduced integration efficiency in CypA-expressing cells (Fig. 3E), an effect that mirrored the inhibitory effect of this CypA inhibitor on HIV-1 infectivity (Fig. 3A). Since, the efficiency of nuclear entry in the CsA-treated cells was unchanged (Fig. 3E), these results further supported a direct link between CypA-CA interaction and HIV-1 integration, an effect seems to be independent of reverse transcription and nuclear entry block. Similarly, the integration efficiency of G89V and P90A mutants was markedly lower in CypA-expressing cells (Fig. 4F). Even though the efficiency of nuclear entry of these mutants was also reduced in these cells, the level of reduction in integration efficiency was markedly lower than the nuclear entry efficiency (Fig. 4F). Notably, a positive effect of CypA-CA binding on integration is also supported by a report showing that both the CA mutants were blocked at the integration step in primary CD4+ T cells [68], even though neither efficiency of nuclear entry nor integration was calculated. Nevertheless, our quantitative analysis emphasized that HIV-1 CA mutants G89V and P90A are blocked at both the nuclear entry and integration step. Finally, CypA’s effect on HIV-1 DNA integration was not influenced by TRIM5α (Fig. 2), the cellular restriction factor that binds to viral capsid. Collectively, these studies strongly supported the model that CypA-CA interaction regulates HIV-1 integration and this post-nuclear entry effect is most likely independent of this host factor’s known effects on viral DNA synthesis and nuclear entry.

To further strengthen our cellular, molecular, genetic and pharmacologic studies (Figs 1-4) supporting a direct role of CypA in HIV-1 DNA integration, we studied the HIV-1 PIC-the biochemical machinery that carries out viral DNA integration into the host genome in an infected cell [24]. The PIC-associated integrase provides the enzymatic activity required for inserting the HIV-1 DNA into the host genomic DNA [24]. Most importantly, functional PICs that retain the DNA integration activity can be extracted from acutely infected cells for biochemical studies [94, 108, 109]. Therefore, we measured the integration activity of PICs extracted from CypA-depleted cells and compared those to CypA-expressing cells (Fig. 5). The integration activity of PICs extracted from the parental CypA-expressing cells efficiently integrated the viral DNA into the target DNA (Fig. 5A). Notably, the activity of PICs from CypA-depleted cells was significantly reduced (Fig. 5B). Furthermore, calculation of the specific activity of these PICs clearly demonstrated that the lower activity of PICs from the cells lacking CypA was not due to lower PIC-associated viral DNA (Fig. 5C). Additionally, this reduction in PIC activity was independent of the presence or absence of TRIM5α protein, since both PIC-mediated viral DNA integration and specific PIC activity remained lower in cells depleted of both CypA and TRIM5α (Fig. 6). Finally, to pin-point a direct role of CypA on PIC-mediated viral DNA integration, we added recombinant CypA protein to the PICs extracted from both CypA-expressing and CypA-depleted cells (Fig. 7). Interestingly, CypA protein markedly stimulated the activity of PICs extracted from both CypA-expressing and CypA-lacking cells (Fig. 7A-B). Surprisingly, the extent of increase in integration activity in the presence of exogenous CypA protein was markedly higher for PICs extracted from CypA-depleted cells (Fig. 7C). These key results suggest that PIC activity is directly stimulated by CypA and optimal integration activity of PICs is regulated by a threshold level of CypA in this viral complex. Future biochemical studies are required to quantitatively measure the stoichiometric link between CypA levels and PIC activity as well as the exact mechanism by which CypA levels stimulate integration.

Finally, we propose a model supporting a hypothetical mechanism by which CypA could stimulate HIV-1 DNA integration (Fig. 8). We predict that CypA promotes HIV-1 integration by specifically binding to the CA-associated with the nuclear viral replication complex. The premise for this model is derived from an emerging functional link between HIV-1 CA and post-nuclear entry steps of virus infection. Particularly, recent reports suggest that an operational intact or near intact capsid lattice enters the nucleus of an infected cell [90–92]. There is also strong evidence that HIV-1 CA is associated with viral replication complexes (RTC/PIC) in the nucleus [20, 22, 112, 113] and is required for the binding of host factors for HIV-1 DNA targeting [114, 115]. Finally, our laboratory has reported that HIV-1 CA is associated with the functional PICs and the levels of PIC-associated CA is directly linked to HIV-1 DNA integration [94]. Therefore, it is plausible that CypA binds to the CA-associated with the viral replication complex (RTC/PIC) in the nucleus and facilitates optimal viral DNA integration into the host chromosomes. Although speculative, we predict that CypA promotes HIV-1 integration by regulating the nuclear CA uncoating process and by extension the optimal engagement of the PIC with the host chromatin. Accordingly, lack of CypA disrupts these CypA-dependent intranuclear processes, resulting in reduced viral DNA integration. Finally, emerging evidence (presented in the 2024 Retroviruses meeting) suggests that CypA likely protects the virus from the inhibitory binding of cytosolic CPSF6 to the virus and that CypA is stripped off of the capsid during passage through the NPC. Therefore, it is possible that a residual amount of CypA remains associated with the HIV-1 CA lattice after nuclear entry and this nuclear CypA promotes PIC-associated viral DNA integration.

**Figure 8.**
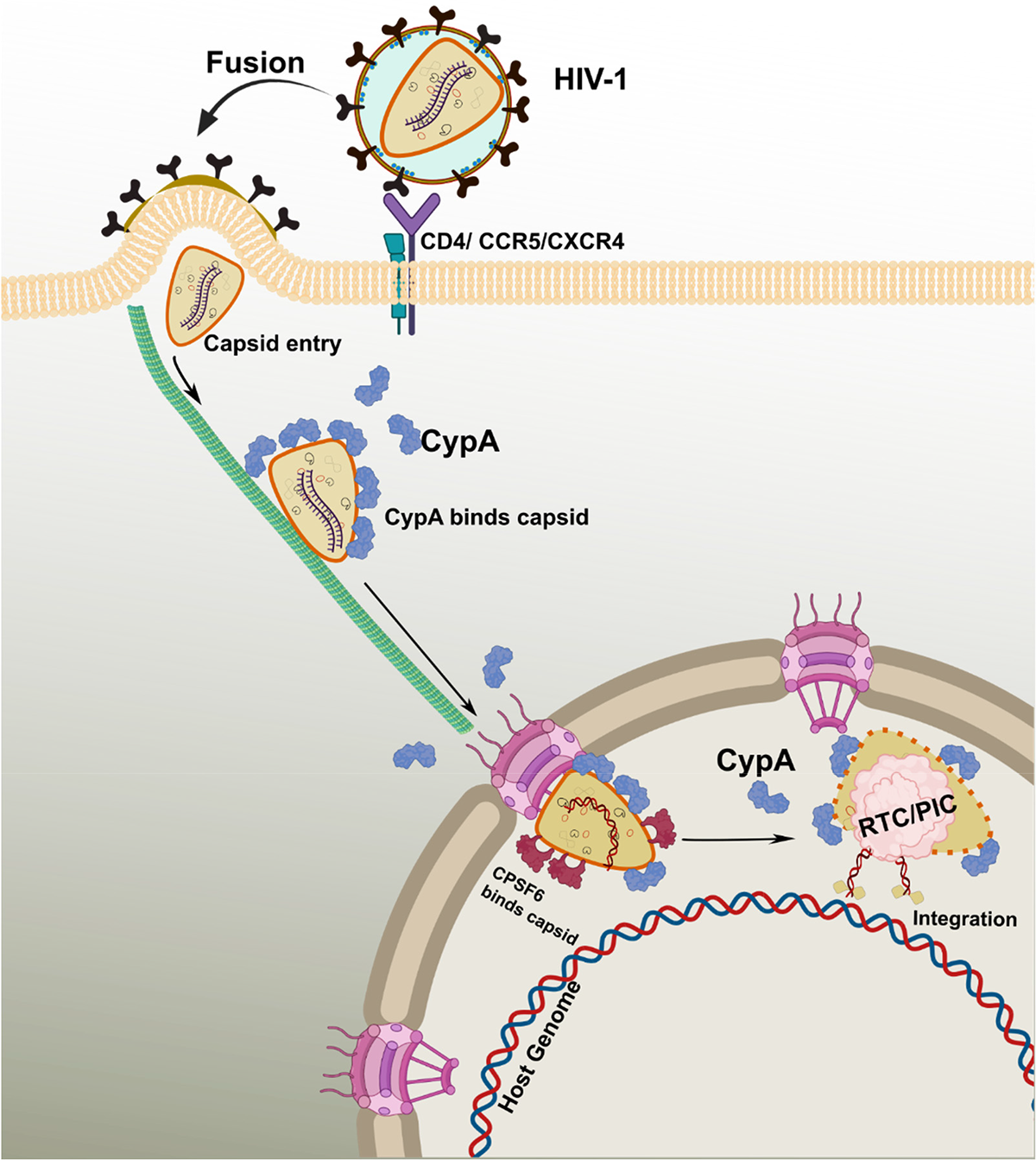
A hypothetical model supporting the mechanism by which CypA promotes HIV-1 DNA integration. The HIV-1 envelope spike binds to the CD^4+^ receptor and one of the co-receptors-CCR5 or CXCR4. The binding leads to the fusion of the viral and target cell membranes and the release of the capsid into the cytosol. HIV-1 capsid transported through microtubules (shown as green elongated tube) where cytosolic CypA (deep blue colored structures) binds to the capsid shell. En route to the nucleus, the operational intact capsid containing the reverse transcribed viral DNA enter the nucleus through the NPC (magenta) and binding with other host factor such as CPSF6 as shown in deep red structures. Upon nuclear entry, CypA remains bound to the near intact capsid and then associated with viral replication complex (RTC/PIC) as shown in light pink blob at site of the integration. The capsid bound CypA in the nucleus promotes the integration of the viral DNA into the host genomic DNA.

## ACKNOWLEDGEMENTS

This work is partly supported by National Institutes of Health grants R01DA057204, R01AI170228, R01AI162694, R01AI136740, and R25AI164610 to CD. This work is also in part supported by the Research Centers in Minority Institutions (RCMI) grant U54MD007586 and the Tennessee CFAR Grant P30AI110527.

